# Aging at scale: Younger dogs and larger breeds from the Dog Aging Project show accelerated epigenetic aging

**DOI:** 10.1101/2024.10.03.616519

**Authors:** Brianah M. McCoy, Blaise L. Mariner, Claire F. Cheng, Elizabeth Slikas, Christine Adjangba, Ashlee Greenier, Layla Brassington, Abbey Marye, Benjamin R. Harrison, Maria Partida-Aguilar, Tal Bamberger, Yadid Algavi, Efrat Muller, Adam Harris, Emily Rout, The Dog Aging Project Consortium, Anne Avery, Elhanan Borenstein, Daniel Promislow, Noah Snyder-Mackler

## Abstract

Dogs exhibit striking within-species variability in lifespan, with smaller breeds often living more than twice as long as larger breeds. This longevity discrepancy also extends to health and aging–larger dogs show higher rates of age-related diseases. Despite this well-established phenomenon, we still know little about the biomarkers and molecular mechanisms that might underlie breed differences in aging and survival. To address this gap, we generated an epigenetic clock using DNA methylation from over 3 million CpG sites in a deeply phenotyped cohort of 864 companion dogs from the Dog Aging Project, including some dogs sampled annually for 2-3 years. We found that the largest breed size tends to have epigenomes that are, on average, 0.37 years older per chronological year compared to the smallest breed size. We also found that higher residual epigenetic age was significantly associated with increased mortality risk, with dogs experiencing a 34% higher risk of death for each year increase in residual epigenetic age. These findings not only broaden our understanding of how aging manifests within a diverse species but also highlight the significant role that demographic factors play in modulating the biological mechanisms underlying aging. Additionally, they highlight the utility of DNA methylation as both a biomarker for healthspan-extending interventions, a mortality predictor, and a mechanism for understanding inter-individual variation in aging in dogs.

## Introduction

Aging is a universal process that varies significantly across and within species. Dogs, in particular, present a unique opportunity to study intraspecific variation in aging due to their remarkable diversity in breed size, lifespan, and other demographic factors [1,2]. Unlike the general trend observed across mammalian taxa, where larger animals typically live longer, larger individuals often have shorter lifespans than their smaller counterparts within species—such as in dogs and many other mammals [3]. Dogs, in particular, present an extreme example of this phenomenon, with larger breeds having significantly shorter lifespans than smaller breeds [4–6]. For instance, larger breeds like Great Danes have an average lifespan of 6-8 years, whereas smaller breeds like Chihuahuas can live up to 15-20 years [7,8]. This inverse relationship has sparked considerable interest in understanding the underlying biological mechanisms that drive aging and longevity in dogs. In addition to breed size, sex, sterilization, and genetic background (purebred vs. mixed breed) also impact longevity and age-associated diseases in dogs [6,9,10].

In addition to size, studies of aging in dogs can dissect the relative impact of other factors on health and aging: (1) Like most mammals, female dogs generally live longer than males, a difference often attributed to the protective effects of sex hormones like estrogen, which may lower the risk of age-related diseases and the potentially harmful impact of the male Y chromosome [11–14]. (2) Sterilization status also influences health trajectories in dogs, with intact dogs showing differential risks for various diseases, such as cancers and reproductive problems, compared to their spayed/neutered counterparts [15,16]. In contrast, sterilized dogs have been shown to have increased risks for conditions like obesity and orthopedic problems [17,18]. (3) Purebred dogs, known for their higher susceptibility to genetic disorders and increased rates of various diseases compared to mixed-breed dogs, tend to have shorter lifespans [19–21]. This disparity is largely attributed to the reduced genetic diversity in purebred dogs, which can lead to an increased prevalence of breed-specific health issues and a greater likelihood of inheriting deleterious alleles [22]. Yet we still know little about how these factors pattern aging at the molecular level, which will provide tremendous insight into the progression of aging and age-related diseases within dogs with immediate relevance to human health.

In addition to “intrinsic” factors (genetics, sex), extrinsic factors such as environmental contaminant exposure and diet significantly influence dog aging trajectories and overall health [23,24]. In this regard, the companion dog is a particularly good model for studying the influence of environmental factors on human aging, as they share their environments [25]. Socioeconomic status, lifestyle choices, and access to healthcare impact health outcomes in humans and dogs, allowing for a comparative analysis of how these factors influence aging across species [20,26–28]. For example, studies have used dogs to model human cardiovascular disease, showing how diet and exercise influence heart health [29,30]. Other research has explored the use of dogs as a model for the impact of environmental toxins on canine cancer rates, providing insights relevant to human cancer epidemiology [31,32]. Additionally, studies on canine cognitive dysfunction (CCD) provide valuable data on aging and neurodegeneration, as CCD shares many parallels with Alzheimer’s Disease. [33,34].

Efforts to characterize inter-individual differences in aging have focused on identifying individual or composite biomarkers that capture an individual’s “biological” age, which can differ from their chronological age [35,36]. Among these biomarkers, DNA methylation (DNAm) predominates, offering insights into the dynamic interplay between genes, the environment, and aging. Changes in DNAm patterns with chronological age have been used to estimate “biological” aging, which can differ from chronological age. Biological age is a biomarker of aging that may be more closely linked to aging outcomes, such as functional decline or mortality, than chronological age [37–40]. Age-associated DNAm changes, often termed “epigenetic aging,” have led to the development and application of DNAm-based biomarkers (“DNAm clocks”) in humans and other species [37,41–44].

To date, researchers have developed a few canine epigenetic clocks to test for breed or species differences in biological aging [45–47]. Still, these efforts are limited by small sample sizes and/or reliance on array technologies that capture only a limited number of highly conserved CpG sites across mammals [45,47]. Here, we extend DNAm clocks to a large and diverse cohort of companion dogs from the Dog Aging Project to understand how extrinsic vs. intrinsic factors influence aging. We leverage our dataset of hundreds of dogs to develop a novel DNAm clock that allows us to (i) prospectively assess changes in biological age over time within dogs and (ii) investigate how demographic factors predict differences in chronological and biological age in companion dogs. Using samples from 864 dogs–448 of which were sampled across multiple years–we first aimed to establish the reliability of the DNAm clock as a biomarker of aging and further hypothesized that it would predict future mortality in our cohort.

Using this new DNAm clock, we tested whether individual attributes, like breed, size, and sex, were linked to accelerated (or decelerated) epigenetic aging. We tested three specific hypotheses: (1) Given the well-established findings that larger dogs have shorter lifespans, we tested if larger breeds exhibit more advanced biological ages than smaller breeds. (2) Since males in many species, including dogs, often age more rapidly than females, we tested if males show accelerated biological aging relative to females [10]. (3) Because purebred dogs have higher rates of many breed-specific diseases [48], we tested if purebred dogs epigenetically aged more quickly than mixed-breed dogs. (4) Lastly, because it has been reported that there are differential effects of sterilization status on health, we tested if sterilization status modifies epigenetic aging in our cohort.

Our clock accurately predicted biological age, highlighting its translatability across diverse breeds and life stages. Additionally, we used the clock to examine how demographic factors—such as breed size, sex, and life stage—may influence the rate of aging in companion dogs. This study reinforces the value of dogs as models for human aging, providing meaningful cross-species insights into the aging process. The development of a more generalizable DNAm clock enables future longitudinal studies to investigate these associations further and understand how these factors affect aging in more diverse cohorts, which will guide the development of targeted interventions to promote healthy aging in dogs.

## Results

### Study cohort

Our sample consisted of 864 dogs in the Precision Cohort (n=420 male, n=444 female; **Figure 1, SI Table 1**) that are a part of the Dog Aging Project [49,50]. The cohort captured most of the canine lifespan (median age: 4.4 years; range: 8 months - 18.5 years; **Figure 1A, SI Table 1**), including 117 recognized pure breeds with representation from the major breeds owned in the US (**Figure 1B, SI Table 2**), and the full range size classes (**Figure 1C**). The dogs also represent different life stages (Puppy: 91, Young adult: 367, Mature adult: 378, Senior: 28, **SI Table 1**).

**Figure 1.**
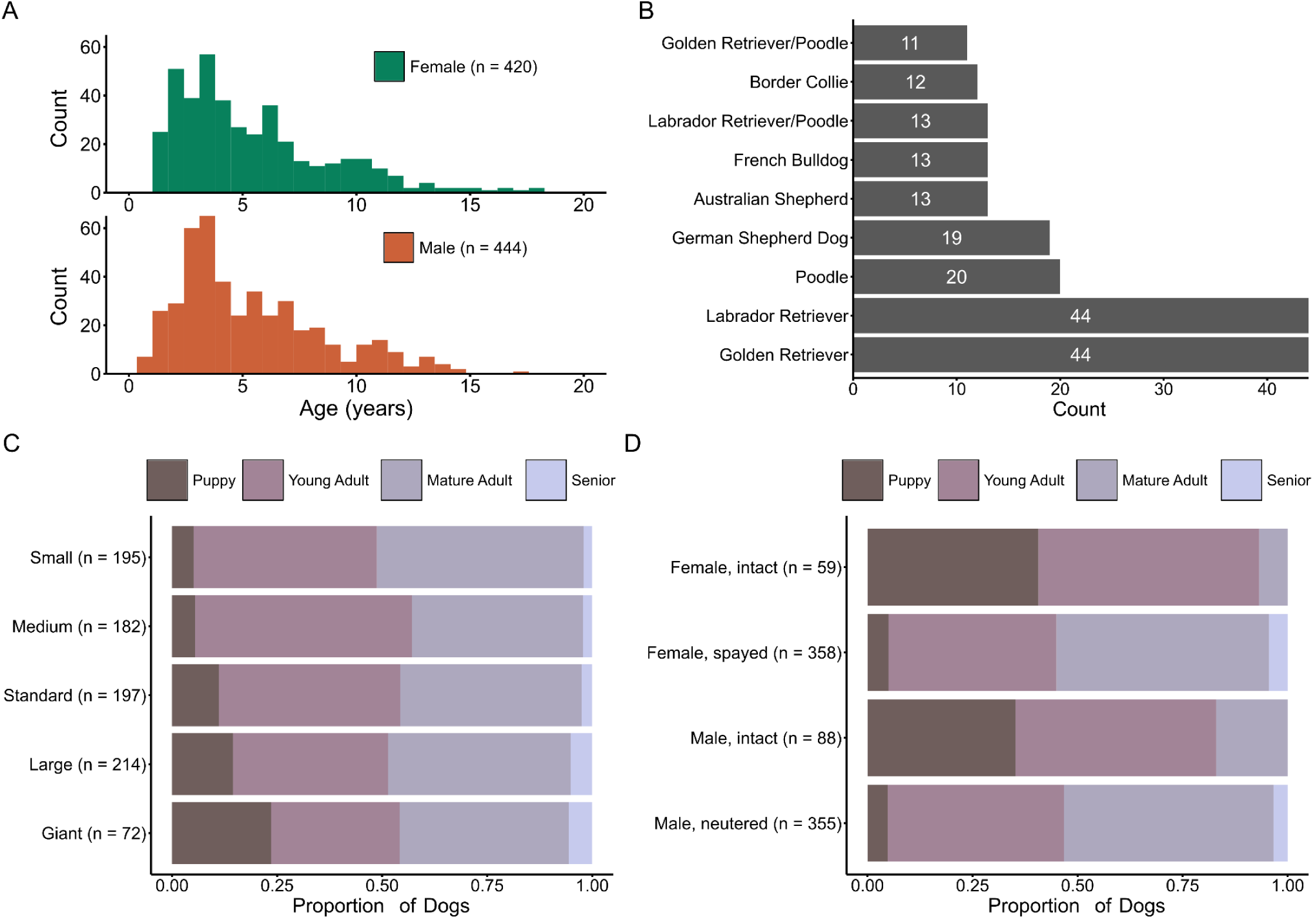
Demographic characteristics of the study cohort. (a) Histograms of age distribution of male and female dogs in the present study. (b) Top 10 most represented breeds in the present study. (c, d) The proportion of dogs in different life stage categories (puppy, young adult, mature adult, senior) stratified by (c) breed size class and (d) reproductive status (data provided in **SI Table 1**).

### Cross-sectional evaluation of DNAm clocks

We used Reduced Representation Bisulfite Sequencing (RRBS) to quantify DNAm at 3,063,134 CpG sites (methods, **Supplemental Figure 1**). We reduced the dimensionality of these 3+ million sites, and thus our multiple hypothesis testing burden, by pooling largely collinear methylation information across nearby CpG sites using a density-based approach where CpG sites within 250 bp of another CpG site were clustered together into 47,393 CpG-dense regions [51]. We then calculated percent methylation for each region for each sample and developed a DNAm clock using elastic net regression with leave-one-out cross-validation (LOOCV; **Figure 2A**, methods).

**Figure 2.**
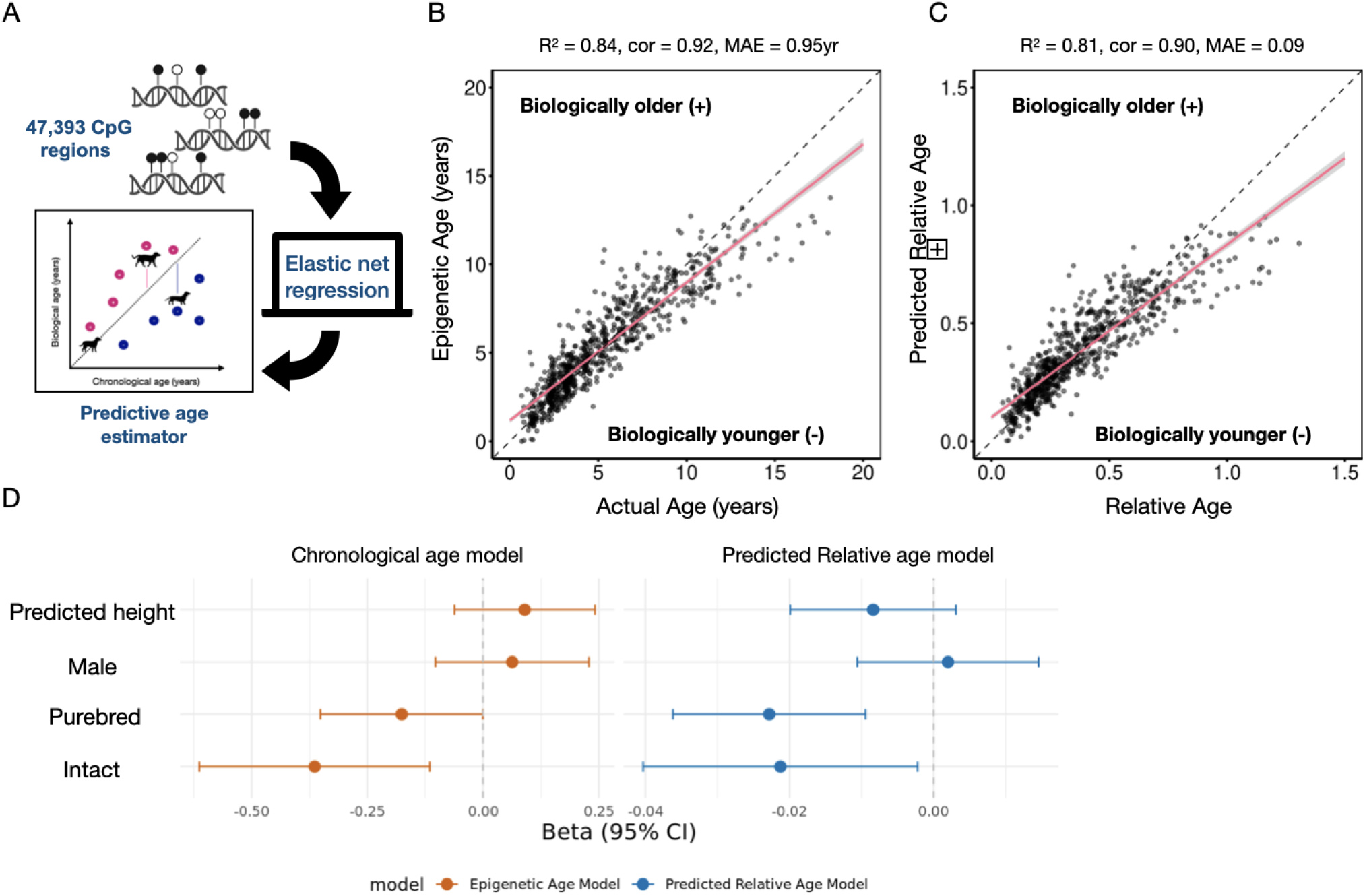
Epigenetic aging in the precision cohort. (a) CpG-dense regions were used to build epigenetic clocks (for region generation overview, see Supplemental Figure 1). (b) Chronological age DNAm clock, predicting age in years. (c) Relative age DNAm clock, predicting biological age relative to breed-specific lifespan (i.e., the proportion of their total expected lifespan that they have lived). (d) Model comparison for chronological and relative age vs. demographic associations.

We generated two measures of DNAm age: one for predicting chronological age (in years) and another for predicting *relative* age, defined as the estimated proportion of the maximum life expectancy of a dog that age. To estimate each dog’s life expectancy, we used the maximum lifespan for their breed, estimated from genetic sequencing and ancestry data (methods, **Figure 2B & 2C**). This approach is similar to that used in another study to increase the translatability of dog DNAm clocks across breeds [45]. We used both chronological and relative age clocks to test whether the rate of aging scales with life expectancy across different dog breeds. For example, Toy Poodles typically have a lifespan of 15 years, while Great Danes live around 8 years. If larger breeds like Great Danes age more quickly than smaller breeds, we would expect a steeper slope in the chronological age clock for the larger breeds, indicating “faster” biological aging. If the relative age clock shows similar slopes across breeds, it suggests that aging is proportional to each breed’s lifespan. By comparing the slopes of these two clocks across breeds with different life expectancies, we can determine how aging processes vary relative to life expectancy.

Our chronological age clock predicted age with an accuracy of R^2^ = 0.85 and a mean absolute error (MAE) of 0.95 yrs (**Figure 2A**), whereas our relative age clock achieved an R^2^ = 0.81 and an MAE = 0.08 proportion of maximum life expectancy (**Figure 2C**). While accurate overall, our chronological age clock varied in how well it predicted chronological age across the lifespan (Figure 2B). The model often predicted younger dogs to be older than their chronological age, while older dogs were predicted to be biologically younger than their chronological age. This underprediction likely reflects a “survivor bias” in our cohort where the sampled dogs have to be alive, which means that the older dogs sampled in our cohort likely reflect “healthier” older dogs that have not yet died. To address this issue, we modeled epigenetic age as a function of our demographic variables while including chronological age as a covariate. This corrects for the bias in predictions across the chronological age spectrum and allows for the effects of other covariates to be interpreted as influences on age acceleration *relative to the expectation based on chronological age* without being confounded by the non-linear association between epigenetic and chronological age. This modeling approach was recently shown to exhibit the least amount of bias in testing for age acceleration using DNAm clocks [52].

To test whether our chronological age clock could predict mortality in our cohort, we first identified dogs that had died after their first year of sampling (n = 49). We then used a Cox proportional hazards model to assess the relationship between residual epigenetic age and mortality. We found that residual epigenetic age was significantly associated with mortality risk (β = 0.29, p = 0.014). In other words, for each unit increase in residual epigenetic age, the risk of death increased by approximately 34% (HR = 1.34, 95% CI: 1.06–1.69). These findings provide validation that our clock is capturing some meaningful variation in biological aging by showing that dogs with higher residual epigenetic age, or those aging more rapidly than expected for their chronological age, have an elevated mortality risk.

We then used a linear model to test for epigenetic age associations with sex, breed status, predicted height, and spay/neuter status while controlling for chronological age [53]. To estimate adult dog size, we included a predicted height metric using a highly predictive random forest model trained in another study of thousands of dogs (see methods). Of the demographic variables tested, we found that spay/neuter status was significantly associated with epigenetic age, with intact dogs being epigenetically younger than their spayed or neutered counterparts (β_intact_ = -0.36, p = 0.004). Interestingly, sex was not a significant predictor of epigenetic age in this model, and neither predicted height nor breed status showed statistically significant associations with epigenetic age (**SI Table 3**). Given that spay/neuter status, but not sex, was significantly associated with epigenetic age, we further investigated whether the effect of spay/neuter status on epigenetic age differed between sexes. To test this, we categorized dogs into four groups: intact males, intact females, neutered males, and spayed females. We found that intact male and female dogs had younger epigenetic ages than their spayed/neutered counterparts (β_intact male_= -0.37, p = 0.008; β_intact female_ = -0.64, p = 0.0005, **SI Table 3**).

Given that most owners sterilize their dogs within the first year of life, many of our intact dogs were on the younger end of our age distribution. Despite this, the lower epigenetic ages of intact dogs suggest that the impact of spay/neuter status on aging is not solely due to age differences. After age-matching the groups, the effect sizes remained consistent, indicating that sterilization likely has a biological influence on epigenetic aging, although the results were no longer statistically significant due to reduced sample size (**SI Table 3**).

This cohort of the DAP, referred across the DAP as the Precision Cohort, is intentionally diverse. We, therefore, also deployed a relative age clock, which adjusts for age differences in life expectancy among breeds, as it may have improved statistical power for testing the effects of demographic factors among a diverse cohort compared to a chronological age clock. Similar to the chronological age clock findings, intact dogs had lower relative ages than spayed/neutered dogs (β = -0.0213, p = 0.029), suggesting that spay/neuter status is linked to aging relative to a dog’s absolute and expected lifespan. The relative age clock also revealed that purebred dogs had lower relative ages than mixed breeds (β = -0.0228, p = 0.0009), suggesting that purebred dogs tend to be biologically younger relative to their expected lifespan. While this effect was statistically significant in the relative age clock, the effect was in the same direction and approached significance in the chronological age clock (**Figure 2D**). In fact, overall, our chronological and relative age clocks revealed concordant demographic effects on epigenetic age acceleration (**SI Table 4; Figure 2D**). Given this result and that the relative age clock requires genetic ancestry data that might not be available in other studies, we chose to focus on the chronological age clock for the remaining analyses to ensure greater translatability to other datasets and ease of interpretation and consistency with existing literature.

### Longitudinal sampling reveals variations in the pace of epigenetic aging across the lifespan

Our longitudinal study design allowed us to test what factors predicted within-individual changes in epigenetic age. Specifically, we collected longitudinal samples from most study dogs 1 year after their first sample (mean = 1.1 years +/-0.072 SD years; see methods). At the time of publication, we have generated longitudinal DNAm data for 52% of our dogs (n = 448), 15% of which we have completed three consecutive years of sampling. We used our chronological age clock (**Figure 2B**) to predict the ages of dogs in these subsequent samples (**Figure 3A**). To estimate the pace of epigenetic aging for each dog, we use the ratio (Δ epigenetic age - Δ actual age), which will be less than zero for dogs that age slowly and greater than zero for dogs aging more quickly and follow this ratio over the longitudinal samples from each dog (**Figure 3A & 3B**). The average aging trajectory (Δ epigenetic age - Δ actual age) of our 437 dogs with longitudinal samples is -0.21 (mean) ± 1.0 year (standard deviation) (**Figure 3B**). To examine potential demographic modifiers of the rate of epigenetic aging, we used a mixed-effects model, with each dog as a random effect to account for repeated measures (67 of 437 dogs had three years of sampling, and thus two epigenetic age differences that we could model). The model tested the difference of change in epigenetic age to change in actual age (Δ epigenetic age - Δ actual age) as a function of sex, age, breed status, reproductive status, and predicted height (**SI Table 5**). Chronological age was the only significant predictor of the rate of aging within individuals (β_age_ = -0.04, p =0.017), suggesting that younger dogs show faster changes in biological age per year lived than older dogs. Overall, we found that 41% of dogs had a faster rate of epigenetic aging compared to their actual age increase (**Figure 3B**), with puppies twice as likely to fall into this accelerated aging category than expected by chance (log_2_(odds ratio) = 2.0, p = 0.03) (**Figure 3C**). These longitudinal results indicate that epigenetic aging is faster in younger dogs.

**Figure 3.**
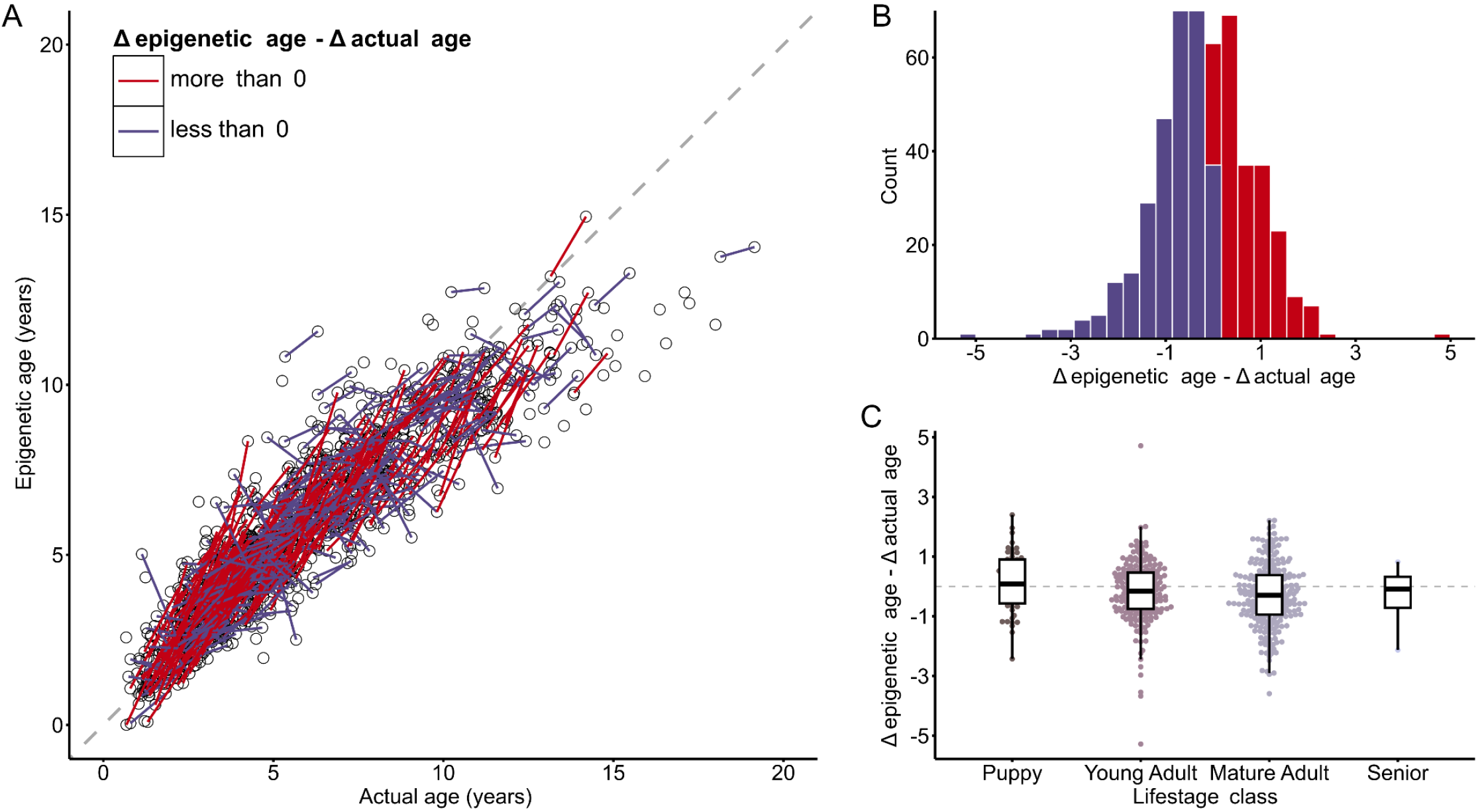
Epigenetic aging in the precision cohort. a) The predicted age of 437 subsequent samples, using the chronological age clock from (2b) (red: change in epigenetic aging > change in chronological age; purple: change in epigenetic aging < change in actual age. b) A histogram showing the distribution of the rates of epigenetic aging. c) Rate of epigenetic aging in different dog life stages (SI Table 5).

### Breed size and sex alter predicted biological aging in cross-sectional epigenetic clocks

To compare differences in epigenetic aging across demographic groups more thoroughly, we also constructed leave-one-demographic-out epigenetic clocks in our cohort. Using the leave-one-demographic-out (LODO) method, we systematically trained epigenetic clocks by excluding specific demographic groups—such as one breed size class or sex—from the training set and testing the model on the excluded group. This approach allowed us to isolate and evaluate how each demographic factor was linked to epigenetic aging by contrasting the relative epigenetic age predictions for excluded groups against those in the training set.

#### Sex

We trained two separate epigenetic clocks: one excluding male data and the other excluding female data during model training. Each clock was then used to predict the epigenetic age of the opposite sex to reveal potential differences in biological aging between males and females [54]. Using this approach, we found that males were predicted to be epigenetically older, on average, than females–suggesting that males age more quickly than females (β_male_ = 0.44, p=2.44e-07; **Figure 4A**). These results align with broader research on vertebrates, where males tend to have shorter lifespans than females, often attributed to sex hormones and/or sex-specific genetic factors. Note that our models do not include CpG sites on sex chromosomes, which are known to contribute to sex-specific aging differences in mice and humans [12,55,56]. This suggests that factors beyond the sex chromosomes, such as hormonal influences or social and environmental experiences, may contribute to the observed differences in epigenetic aging between males and females.

**Figure 4:**
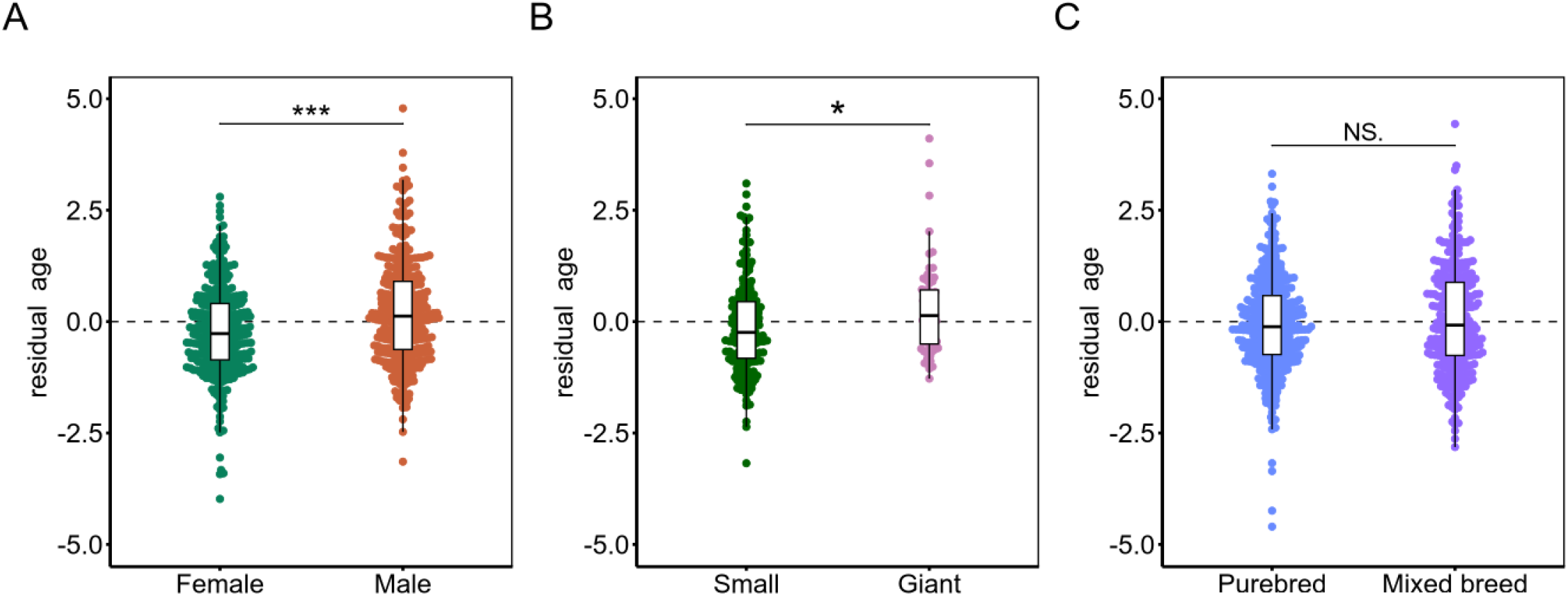
Cross-sectional, leave-demographic-out clock differences in epigenetic aging between sex and breed size. a) Residuals (predicted age minus actual age) from a male-trained clock tested with female samples (red) and a female-trained clock tested with male samples (orange). b) Residuals (predicted age minus actual age) from a non-small-trained clock tested with small dog samples (green) and a non-giant-trained clock tested with giant samples (pink). c) Residuals (predicted age minus actual age) from a purebred-trained clock tested with mixed breed samples (purple) and mixed breed-trained clock tested with purebred samples (blue). All p-values are from linear models (*p<0.05, **p<0.01, ***p<0.001).

#### Breed size

To explore the differences in epigenetic aging between larger and smaller breeds, we developed two leave-breed-size-out epigenetic clocks, following a similar approach to the leave-sex-out clocks, to examine how breed size might affect the rate of epigenetic aging. We found that the model trained on non-small breeds predicted small dogs to be epigenetically younger on average than giant dogs predicted by the non-giant-trained model (β_giant_ = 0.37, p=2.2e-2; **Figure 4B**). This suggests that age-related DNA methylation (DNAm) patterns are accelerated in giant breeds compared to small breeds, potentially contributing to their
shorter lifespans [7,8,48,57].

#### Mixed vs. Purebred dogs

Lastly, we developed two leave-breed-status-out epigenetic clocks to examine whether the shorter lifespans of purebred dogs are associated with accelerated epigenetic aging, compared to their mixed-breed counterparts. Contrary to our analysis of the overall cross-sectional clock, we did not find any significant difference in epigenetic aging between pure and mixed-breed dogs, although the effect size was in a similar direction as the relative age clock showing that purebred dogs were relatively epigenetically younger than their mixed breed counterparts (β_Purebred_ = -0.14, p=0.11; **Figure 4C**).

## Discussion

Here, we present a novel biological age estimator using DNAm profiles from the largest, most diverse cohort of companion dogs. Our novel dataset and clock, which was paired with demographic and genetic data collected from the same dogs, allowed us to identify important individual factors that pattern biological aging in companion dogs–an emerging and exciting model for human health and aging. Leveraging our longitudinal samping and follow-up of this DAP cohort revealed that our clock predicted future mortality above and beyond that of just chronological age, suggesting that it is capturing meaningful variation in biological aging and meets some criteria for being a biomarker of aging. Our longitudinal sampling allowed us to identify what factors predict within-individual changes in aging, often referred to as the “pace of aging”–something that has thus far been absent from most other studies outside of those in humans and lab organisms. Finally, by developing and deploying this clock in our large and diverse cohort, we were able to make four key findings that advance our understanding of variation in aging in dogs that may translate to ourselves.

First, our DNAm clock accurately predicts chronological and relative ages across diverse dog breeds. Other studies have developed predictors of age in dogs, beginning with Thompson et al. (2017) and later by Horvath et al. (2022). Unlike these earlier efforts that used array-based technology focusing on single CpG sites, we constructed our epigenetic clock using a density-based region approach. This method offers two advantages: (i) it significantly narrows the search space, reducing complexity by examining discrete regions of CpGs instead of millions of individual sites (e.g., we covered about 3 million CpGs post-filtering). (ii) regions each capture multiple CpGs, providing more robust estimates for downstream analysis by encompassing multiple CpGs, offering a more comprehensive view of methylation patterns and reducing variability compared to single sites and coverage variability often seen in Reduced Representation Bisulfite Sequencing (RRBS) data. This approach enhances the reproducibility and applicability of methylation-based aging models across different datasets, making them an attractive tool for predicting biological age in diverse dog populations. In contrast to previous canine epigenetic aging models, our chronological age-based model addresses limitations such as sample size and demographic diversity. Models constructed using data from a narrow range of breeds–predominantly purebred dogs–restrict their applicability across the diverse samples. Our model, in contrast, incorporates a cohort of 864 dogs that includes over 100 breeds and a range of mixed breeds, enhancing generalizability in predicting biological age. This comprehensive approach provides a reliable tool for studying the complex dynamics of aging in a broader canine population, which we used to test factors that may modify the aging process in dogs.

Second, we demonstrate that the chronological age clock can effectively predict mortality in our canine cohort, highlighting its utility beyond simply estimating biological age. The significant association between residual epigenetic age and mortality risk suggests that dogs with higher-than-expected epigenetic ages relative to their chronological ages are at an increased risk of death. This indicates that our DNAm clock captures underlying biological aging processes relevant to survival outcomes. Future work will explore the potential of DNAm clocks as predictive tools for the healthspan and lifespan of dogs in our cohort. We can refine these models by integrating this epigenetic data with other health metrics to better predict disease onset and mortality. This approach could facilitate the identification of dogs at higher risk for age-related health issues, enabling earlier interventions and personalized care strategies. Additionally, longitudinal sampling and follow-up studies will be used to validate the predictive accuracy of these clocks over time, ultimately contributing to our understanding of aging mechanisms and improving health outcomes for both dogs and humans.

Third, we deployed our clock on longitudinal data, which allowed us to characterize variation in rates of aging among individuals–something that is not possible using cross-sectional data alone. There are inherent limitations to cross-sectional studies of aging–particularly those that capture samples covering a wide range of ages. One limitation is the presence of survival bias, in which older subjects can, by nature, only be the subjects that survived to that age. Indeed, we see evidence of such bias in this cohort, where the epigenetic age of older dogs is lower than expected based on their chronological age (**Figure 2B & C**). Some have argued that cross-sectional analysis is biased against discovering aging biomarkers [58]. Whereas strictly cross-sectional data may reveal differences between young and old subjects, even with great accuracy, such data may fail to identify biomarkers of *aging*.

One way to mitigate this bias is to conduct longitudinal studies by collecting repeated samples of individuals over time. Such studies, however, are expensive and time-consuming and, as such, have not routinely been conducted in dogs. Here, we used follow-up data within the Precision cohort to test the hypothesis that longitudinal follow-up data can reveal demographic factors that can influence the trajectory or rate of aging. We found that the increase between the first and second year of sampling is predicted by our clock (0.90 ± 0.05 years, **Figure 3A & 3B**) and that the age increase over this interval is not modified by either sex, or dog size, with most of the Δ epigenetic age - Δ actual age differences likely explained by the dogs’ life stage (**Figure 3C**). This result suggests that neither size nor sex significantly influences the rate dogs age epigenetically. However this could be due to only having follow-up DNAm data from half of our cohort. We are currently generating additional longitudinal data from the rest of the cohort across multiple years to better test this.

Lastly, our analysis using both the cross-sectional and LODO clocks revealed valuable insights into the effects of demographic factors on epigenetic aging. Using our cross-sectional clock, we found evidence of an effect of spay/neuter status on epigenetic aging, particularly in females; there was not a significant difference in female epigenetic age prediction from males. One limitation is that intact dogs in our dataset are generally younger than their spayed/neutered counterparts. This imbalance in age ranges between groups may have masked the more pronounced effects of spay/neuter status on aging. A more balanced dataset, especially with older intact dogs, would be better suited to explore the long-term effects of spay/neuter status on epigenetic aging. Our LODO methods helped isolate the effects of sex and breed size on the aging process. We found male genomes to be epigenetically older than female genomes despite differences in age (**Figure 4A**). These findings align with some research suggesting that female dogs often live longer than males, often explained by differences in hormone levels and sex chromosomes [12,59]. This observed sex-based difference in aging rates mirrors similar findings in human studies, where women generally live longer than men, possibly due to hormonal influences and lifestyle factors [60]. Notably, our model found epigenetic differences in aging by training our models with autosomal genome region information only, indicating a differential loss in epigenetic regulation between sexes on autosomes that has yet to be understood. The LODO clocks showed that larger breeds tend to have an older epigenetic age on average compared to smaller breeds (**Figure 4B**). Although this trend was not statistically significant in the findings from our chronological age clock, the direction of the association was consistent. All of this supports the established knowledge that larger breeds have shorter lifespans and faster aging rates [57]. Although the exact role of epigenetics in the aging process remains unclear, whether it actively contributes to aging or merely reflects it, much of the variation in lifespan between large and small dogs is frequently attributed to differences in growth hormone exposure [61,62]. This opens the door to understanding how these hormonal differences interact with epigenetic mechanisms to produce disparities in aging between breeds.

Contrary to our expectations, the leave-one-demographic-out (LODO) clock did not show significant differences in epigenetic aging between mixed-breed and purebred dogs, although the effect size suggested that purebred dogs were epigenetically younger than mixed-breed dogs. This matches the results of the relative age clock, where purebred dogs had significantly lower relative ages than mixed breeds (β_purebred_ = -0.0228, p = 0.001, **SI Table 4**). Taken together, these results suggest that purebred dogs may age more slowly relative to their expected lifespan despite their overall shorter absolute lifespans. One possible explanation is that the relative age clock may underestimate the life expectancy of mixed-breed dogs because their genetic diversity contributes to a broader range of aging trajectories. In contrast, purebred dogs, with their more controlled genetic backgrounds, might have more consistent aging patterns. Still, these breed-specific traits could also mask differences in epigenetic aging when averaged across a larger population. Finally, the variability in health and environmental factors within each group might further dilute the strength of the association between breed status and epigenetic aging, making it harder to detect a clear difference.

While this study offers new insights into canine aging and validates the impact of demographics on aging, there are a few notable limitations to consider. First, while our cohort is large, it represents only a subset of the diversity within the DAP Pack and the global population of companion animals [49,63]. As a result, some associations between demographics and epigenetic aging were not statistically significant but showed trends in the expected direction. This could be due to a lack of power to detect these effects fully. Expanding the cohort to include a more diverse range of breeds, a larger number of individuals, and repeated sampling could enhance the statistical power and potentially reveal more significant associations.

Additionally, our study relies on some cross-sectional data, which captures a single point in time for each dog. Extending our current two-year longitudinal data to track the same individuals over a longer period would provide a more comprehensive understanding of the aging process. This would allow us to observe changes in our epigenetic biomarkers and their relationship with health outcomes more accurately. Our study is also limited because our current analysis does not include the potential influence of environmental factors, such as diet, exercise, socioeconomic status, and living conditions, as it was outside the scope [64–67]. These factors can significantly impact aging and may confound the observed associations. Future studies will aim to account for these variables to isolate the specific effects of demographic factors on epigenetic aging. Moreover, to better understand the impact of inbreeding and breed-specific health issues on biological aging, it is crucial to increase the sample size to include more dogs from breeds with shorter lifespans (e.g. Great Danes, Dalmatians).

This study lays the foundation for future research by identifying potential causes and consequences of aging while also highlighting the immense potential of the Dog Aging Project as a powerful model for translational, age-related research. As we continue to collect more data and track these dogs throughout their lifespans, we will have the opportunity to determine if and how epigenetic profiles can predict future health trajectories. This will allow us to explore the relationships between environmental risk factors and long-term health outcomes, deepening our understanding of how aging unfolds in companion animals and humans. The unique demographic characteristics of dogs, coupled with longitudinal data collection, will provide valuable insights that can inform strategies to promote healthy aging across species.

## Methods

### Study population

Samples were obtained from The Dog Aging Project (DAP), a community science project aiming to understand how genes, lifestyle, and the environment influence aging and disease outcomes [49,50] For this study, we obtained whole blood, data collected during veterinary clinic visits, and survey data from DAP’s Precision cohort (864 unique dogs: 780 year 1 samples, 496 year 2 samples and 103 year 3 samples).

The University of Washington IRB deemed that recruitment of dog owners for the Dog Aging Project, and the administration and content of the DAP Health and Life Experience Survey (HLES), are human subjects research that qualifies for Category 2 exempt status (IRB ID no. 5988, effective 10/30/2018). All study-related procedures involving privately owned dogs were approved by the Texas A&M University IACUC, under AUP 2021-0316 CAM (effective 12/14/2021).

### Whole genome sequencing, ancestry estimation, and height prediction

The DAP utilized low-pass whole-genome sequencing and imputation to establish ancestry. Sequencing reads were aligned to the CanFam3.1 reference genome (NCBI GCF_000000145.2). Genotype imputation was done using loimpute and the dog low-pass v2.0 [0.1x-6x] panel, representing 540 dogs across ∼133 breeds, 28 mixed-breed dogs, and various Indigenous and wild canines. Public genotype data from 109 modern breeds, 3 village dog populations, and 2 wolf populations were used for ancestry inference. Ancestry-informative markers were identified using PLINK2, and ADMIXTURE in supervised mode was employed to infer ancestry from these populations. Using this genotypic data, we predicted adult heights using a random forest model trained and validated in a study of 1,730 dogs [68] with paired owner-reported height data (scale 0-4: 0: ankle high; 1: calf high; 2: knee high; 3: thigh high; 4: waist high).

### PBMC isolation & DNA/RNA extraction

Peripheral blood mononuclear cells (PBMCs) were isolated from whole blood at the DAP Central Lab at Texas A&M University College of Veterinary Medicine & Biomedical Sciences. We then extracted DNA and RNA from PBMCs from 1379 samples from dogs in the Precision cohort using (Zymo Quick DNA/RNA Magbead Extraction Kit).

### Reduced Representation Bisulfite Sequencing (RRBS) and CpG region generation

Our established protocol generated RRBS libraries from ∼200 ng of DNA [69]. Samples were then sequenced to an average depth of 16.7 million reads per sample on the Illumina NovaSeq platform at the Translational Genomics Research Institute’s Collaborative Sequencing Center. Sequenced reads were trimmed and mapped to the dog genome using Trim Galore! (https://github.com/FelixKrueger/TrimGalore) and bismark2 [70]. For all downstream analysis, the Labrador retriever genome (UU_Cfam_GSD_1.0_ROSY) was used as a reference assembly (GCF_014441875.1)[71].

Our analyses only included CpG sites with≥ 5x coverage across ≥ 33% of samples (n=864 dogs). Each CpG region in the analysis contained at least 5 CpG sites, with adjacent sites separated by no more than 250 base pairs. We then filtered out regions that were consistently methylated (median methylation > 90%) or consistently unmethylated (median methylation < 10%) to focus on those with variable methylation patterns in our cohort, where we have the statistical power to detect age-associated differences in DNAm. This filtering process resulted in 47,393 regions for downstream analysis (**Supplemental Figure 1B**). At the end of region generation and filtering, regions had an average length of 340 (SD +/-336) nucleotides and 16 (SD +/-20) CpG sites, with an average of 24 (median = 21) nucleotides separating CpG sites within each region (**Supplemental Figure 1C**). We then calculated percent methylation for each sample in each region and imputed missing values (1.5% total missing values across all samples and all regions) using the R package methyLImp2 [72].

### DNA methylation clocks

Due to experimental limitations, the resulting percent methylation matrices mentioned above may contain missing information. We used the MICE package in R to impute the missing data using predictive mean matching with default parameters. To build our canine DNA methylation clock, we used an elastic net regression (glmnet package in R) and a leave-one-out-cross validation (LOOCV), where all but one of the samples is used to predict the age of the left-out sample. The model has two parameters, an alpha (α) and a regularization parameter lambda (λ). After testing various alpha values, 0.5 yielded the best model performance. The alpha was set to 0.5 to balance the two parameters equally and ensure optimization of feature selection.

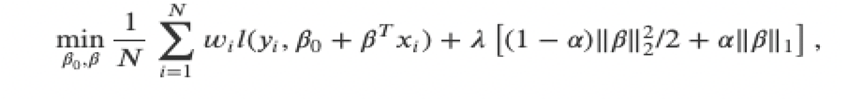

#### Relative age

To allow for the comparison of dog breeds with markedly different lifespans, we used a method adapted from Horvath et al. to generate relative age using the formula relative age = age/max lifespan, where the maximum lifespan for dogs was estimated using the breed determined from genetic sequencing and ancestry data for all dogs in the precision cohort (**SI Table 6**). We took the mean of the reported lifespan ranges from the Kennel Club (KC), the American Kennel Club (AKC), and the Australian National Kennel Council [73–75].

#### Leave-demographic-out

To build the leave-sex-out clocks, dogs not in the current sex category were used as the training set, and dogs in the current sex category were used as the test set. A DNAm clock was trained on the training set using the cv.glmnet () function from the glmnet package, with a 10-fold cross-validation to select the optimal regularization parameter (alpha was set to 0.5). The trained model was then used to predict the ages of the dogs in the test set. The mean squared error (MSE) between the predicted and actual ages and the R^2^ was calculated for each model.

A similar approach was used to construct the leave-one-size-out clocks. Each size category (small, medium, standard, large, & giant) was used as the training set, while the remaining size categories were used as the test set. The glmnet model parameters were the same as those used in the previous models. The mean squared error (MSE) between the predicted and actual ages and the R^2^ was calculated for each model.

## Statistical analysis

All statistical analysis was carried out in R (4.2.2). Linear models of epigenetic age, relative age, average time to death, and breed demographics were fit using ordinary least squares regression with the lm() function in R. Sample size across each model varies due to available metadata (see SI tables for more details).

## Code and data availability

The Dog Aging Project is an open data project. These data are housed on the Terra platform at the Broad Institute of MIT and Harvard.

## Acknowledgments

The authors thank Dog Aging Project participants, their dogs, and community veterinarians for their essential contributions. This research is based on publicly available data collected by the Dog Aging Project, under U19 grant AG057377 (PI: Daniel Promislow) from the National Institute on Aging, a part of the National Institutes of Health, and by additional grants and private donations, including generous support from the Glenn Foundation for Medical Research, the Tiny Foundation Fund at Myriad Canada, the WoodNext Foundation, and the Dog Aging Institute. These data are housed on the Terra platform at the Broad Institute of MIT and Harvard. BMM was supported by an NIA F99/K00 and NSF Graduate Research Fellowship. The authors acknowledge Research Computing at Arizona State University for providing High Performance Computing and Storage resources that have contributed to the research results reported within this paper [76]. The content is solely the authors’ responsibility and does not necessarily represent the official views of the National Institutes of Health.

**Dog Aging Project Consortium Authors (as of March 2022)**

Joshua M. Akey^1^, Brooke Benton^2^, Elhanan Borenstein^3^, Marta G. Castelhano^4,5^, Amanda E. Coleman^6^, Kate E. Creevy^7^, Kyle Crowder^8^, Matthew D. Dunbar^9^, Virginia R. Fajt^10^, Annette L. Fitzpatrick^11^, Unity Jeffery^12^, Erica C Jonlin^13^, Matt Kaeberlein^14^, Elinor K. Karlsson^15^, Kathleen F. Kerr^16^, Jonathan M. Levine^17^, Jing Ma^18^, Robyn L McClelland^19^, Daniel E.L. Promislow^20^, Audrey Ruple^21^, Stephen M. Schwartz^22^, Sandi Shrager^23^, Noah Snyder-Mackler^24^, Katherine Tolbert^25^, Silvan R. Urfer^26^, Benjamin S. Wilfond^27^

^1^ Lewis-Sigler Institute for Integrative Genomics, Princeton University, Princeton, NJ, USA

^2^ Department of Laboratory Medicine and Pathology, University of Washington School of Medicine, Seattle, WA, USA

^3^ Department of Clinical Microbiology and Immunology, Sackler Faculty of Medicine, Tel Aviv University, Tel Aviv, Israel

^4^ Cornell Veterinary Biobank, College of Veterinary Medicine, Cornell University, Ithaca, NY, USA

^5^Department of Clinical Sciences, College of Veterinary Medicine, Cornell University, Ithaca, NY, USA

^6^ Department of Small Animal Medicine and Surgery, College of Veterinary Medicine, University of Georgia, Athens, GA, USA

^6^7Department of Small Animal Clinical Sciences, Texas A&M University College of Veterinary Medicine & Biomedical Sciences, College Station, TX, USA

^8^ Department of Sociology, University of Washington, Seattle, WA, USA

^8^9Center for Studies in Demography and Ecology, University of Washington, Seattle, WA, USA

^10^ Department of Veterinary Physiology and Pharmacology, Texas A&M University College of Veterinary Medicine & Biomedical Sciences, College Station, TX, USA

^11^ Department of Family Medicine, University of Washington, Seattle, WA, USA

^12^ Department of Veterinary Pathobiology, Texas A&M University College of Veterinary Medicine & Biomedical Sciences, College Station, TX, USA

^13^ Department of Laboratory Medicine and Pathology, University of Washington School of Medicine, Seattle, WA, USA

^14^ Department of Laboratory Medicine and Pathology, University of Washington School of Medicine, Seattle, WA, USA

^15^ Bioinformatics and Integrative Biology, University of Massachusetts Chan Medical School, Worcester, MA, USA

^16^ Department of Biostatistics, University of Washington, Seattle, WA, USA

^17^ Department of Small Animal Clinical Sciences, Texas A&M University College of Veterinary Medicine & Biomedical Sciences, College Station, TX, USA

^18^ Division of Public Health Sciences, Fred Hutchinson Cancer Research Center, Seattle, WA, USA

^19^ Department of Biostatistics, University of Washington, Seattle, WA, USA

^20^ Department of Laboratory Medicine and Pathology, University of Washington School of Medicine, Seattle, WA, USA

^21^ Department of Population Health Sciences, Virginia-Maryland College of Veterinary Medicine, Virginia Tech, Blacksburg, VA, USA

^22^ Epidemiology Program, Fred Hutchinson Cancer Research Center, Seattle, WA, USA

^23^ Collaborative Health Studies Coordinating Center, Department of Biostatistics, University of Washington, Seattle, WA, USA

^24^ School of Life Sciences, Arizona State University, Tempe, AZ, USA

^25^ Department of Small Animal Clinical Sciences, Texas A&M University College of Veterinary Medicine & Biomedical Sciences, College Station, TX, USA

^26^ Department of Laboratory Medicine and Pathology, University of Washington School of Medicine, Seattle, WA, USA

^27^ Treuman Katz Center for Pediatric Bioethics, Seattle Children’s Research Institute, Seattle, WA, USA

## Supplemental Figures

**Supplementary Figure 1:**
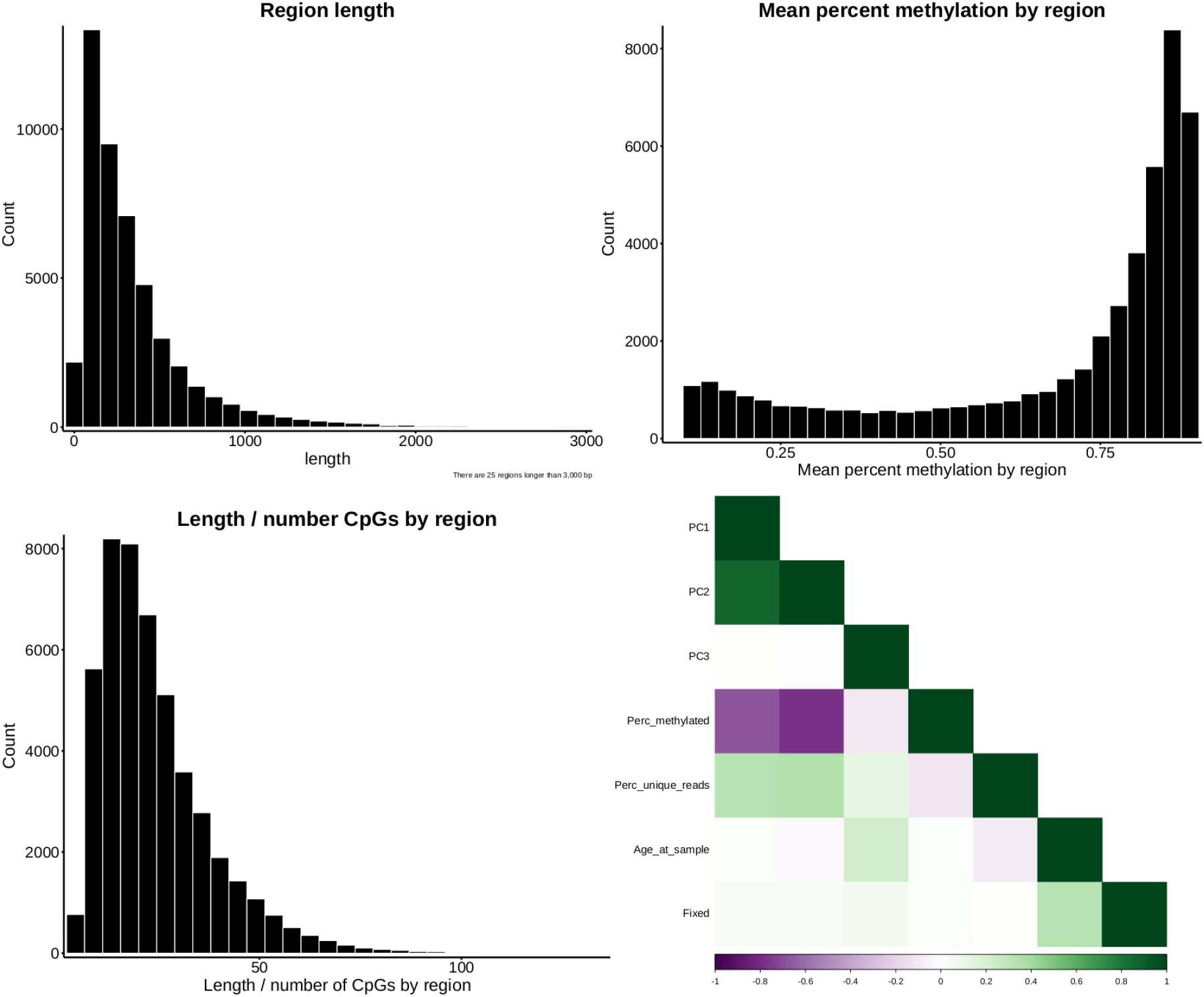
Generation of CpG regions. a) Distribution of region lengths (in bp) with the majority being less than 1000 base pairs. b) Methylation levels across the generated regions, indicating predominantly high methylation. c) CpG site density within the regions. d) Correlation plot showing both technical and biological variance observed in the dataset.

